# A Selenium-Deficient Mouse Model of Mouse-Adapted SARS-CoV-2 Demonstrates Variant Emergence Observed in SARS-CoV-2 Pandemic Variants

**DOI:** 10.64898/2026.08.25.746963

**Authors:** Monica E. Graham, Olukunle O. Oluwasemowo, Deepa Murugesh, Margarita V. Rangel, Jeffrey A. Kimbrel, Aram Avila-Herrera, James Thissen, Adam Zemla, Ashlee M. Phillips, Nicole Collette, Dina R. Weilhammer, Monica K. Borucki

**Author notes:** corresponding author, Biosciences and Biotechnology Division, Lawrence Livermore National Laboratory, 7000 East Ave, Livermore, CA 94551-0808, USA.

## Abstract

Host selenium deficiency has been shown to generate novel genetic variants in RNA viruses. With the predicted rise of selenium deficiency globally, we sought to determine if host selenium deficiency can be a predictive factor for RNA virus variant emergence. We utilized a selenium-deficient BALB/c mouse model to investigate how host selenium status influences the emergence of viral variants in mouse-adapted SARS-CoV-2. Mice were maintained on control or selenium-deficient diets and subjected to sequential rounds of diet-matched viral passage to generate diet-specific virus populations. Deep sequencing of passaged viral populations revealed that selenium-deficient passage drove a marked increase in inter-host genomic heterogeneity and produced a distinct mutational profile relative to control passage. Eighteen mutations were identified as unique to selenium-deficient passage, including variants previously observed during natural human SARS-CoV-2 evolution. These mutations were largely maintained at sub-consensus frequencies, indicating that selenium deficiency can expand the viral quasispecies landscape that enrichs reservoirs of adaptive potential. To determine how this altered mutant spectrum affected pathogenesis, we challenged normal diet-fed adult and aged BALB/c mice with control-or selenium-deficient-passaged virus. Although overt differences in weight loss, survival, and viral burden were generally modest, selenium-deficient-passaged virus induced pronounced increases in antiviral cytokine expression. Together, these findings identify host selenium deficiency as a driver of RNA virus population diversification and show that nutritionally stressed animal models can reproducibly generate mutations observed in nature.

**Importance:** With the increasing global threat of pandemic-potential RNA viruses, it it vital to identify predictive factors and tools to prepare for the next pandemic. Selenium deficiency is an increasingly prevalent form of malnutrition that impacts immune function, oxidative stress, and RNA virus mutagenesis. Based on current data on selenium deficiency and viral variance, we hypothesized that a selenium-deficient mouse model of SARS-CoV-2 infection could generate increased viral diversity as well as recapitulate naturally-occuring variants. Our deep-sequencing data of diet-passaged mouse-adapted SARS-CoV-2 revealed that repeated selenium-deficient passage increased genomic heterogeneity, produced consistent variant profiles across individual mice, and generated human SARS-CoV-2 variants found in nature. Our results demonstrate *in vivo* selenium-deficient viral passage as a potential experimental platform for forecasting putative variant emergence during future viral outbreaks.

## Introduction

Selenium (Se) is a micronutrient obtained through the diet that is critical for multiple facets of human health^1^. In the human body Se predominantly functions in selenoproteins, such as glutathione peroxidase, that play vital roles in controlling oxidative stress and reactive oxygen species generation^1–4^. Se is also crucial for functioning immune responses, including cytokine expression and adaptive immune cell activation^1,3,4^. Human Se intake relies on sufficient Se levels in soil and crops, as Se is obtained mainly through diet via crops and animal products^2,3,5^. Climate change factors such as decreased precipitation and increased soil aridity are predicted to be major drivers of reduced Se soil content^5^. These climate factors, in combination with low Se release from natural sources, are predicted to cause significantly decreased Se content across the globe^5,6^. Approximately 1 in 7 people globally are currently consuming inadequate levels of selenium, and with predicted global decreases in Se soil content, increased prevalence of human Se deficiency may be inevitable^2,5^. Se deficiency significantly increases oxidative stress and alters immune responses, factors that increase severity of viral infections^3,7^. Se deficiency-induced oxidative stress has also been implicated in increased viral genome mutation rates, particularly of RNA viruses^1,4,7^.

Experimental and epidemiological studies have demonstrated associations of host Se deficiency with viral mutagenesis. Researchers have discovered that host Se deficiency during coxsackievirus B3 (CVB3) infection caused increased viral mutation and virulence, including the development of a cardio-virulent CVB3 strain reported in regions China^4,8,9^. Infection of Se-deficient mice with influenza A H3N2 produced similar outcomes of increased viral mutation and symptom severity^4^. Variant emergence is not a random process but a complex interaction between viral, ecological, and host factors contributing to the generation of novel viral variants^1,4,10^. Evidence supports host Se deficiency can lead to increased viral mutation rates and virulence, and climate-influenced Se deficiency may exacerbate these effects^4,5,9,11–13^. With the emergence of novel viruses anticipated to increase with climate change, it is critical to idenfity factors influencing variant emergence that can be utilized for variant emergence prediction.

To this end, we simulated passage of mouse-adapted severe acute respiratory syndrome coronavirus 2 (maSARS-CoV-2) through Se-adequate and Se-deficient populations via a mouse model of diet and infection. We chose maSARS-CoV-2 as our model pathogen due to well-documented sequencing data of human SARS-CoV-2 that would provide a reference for relevant viral variant emergence. Passaged virus populations were analyzed via deep sequencing and variant calling computational tools to evaluate potential changes in variant emergence and genomic heterogeneity. We found that Se-deficiency produced consistent, inter-host increases in viral genomic heterogeneity. We discovered that Se-deficiency not only produced unique variants, but recreated variants that emerged naturally during the COVID-19 pandemic. Most of these novel variants were found at sub-consensus frequencies across assayed mice. We also assessed our Se-adequate and Se-deficient passaged viruses for changes in virulence and pathogenesis in normal diet conditions and found that our Se-deficient virus caused moderate phenotypic changes to maSARS-CoV-2 infection. Together, our findings report that a Se-deficient mouse model of RNA virus infection significantly increases viral mutation frequency and recapitulates human SARS-CoV-2 variants found in nature, suggesting that such a model could be applied to forecast putative variants in emerging RNA viruses.

## Results

### Generating Diet-passaged maSARS-CoV-2 Virus Populations

To model viral spread through nutrient-sufficient and SEDEF populations, we utilized a selenium-deficient mouse model in conjunction with a moused-adapted strain of SARS-CoV-2 (maSARS-CoV-2). Equal numbers of BALB/C mice were fed on a control diet (CTRL) or selenium deficient diet (SEDEF) (Table 1) for four weeks before intranasal infections with 10^5^ PFU maSARS-CoV-2. Lung homogenate was used to passage virus into new cohorts of mice a total of eight times through CTRL-or SEDEF-fed mice. Across mouse passages, percentage weight change only varied between CTRL and SEDEF populations during passage 2 (Figure 1A). MaSARS-CoV-2 titer in pooled lung homogenates did not vary significantly across passages (Figure 1B). Our final virus pools isolated from the eighth passage will be refered to as “P8-CTRL” and “P8-SEDEF” for the remainder of the text.

**Table 1:**
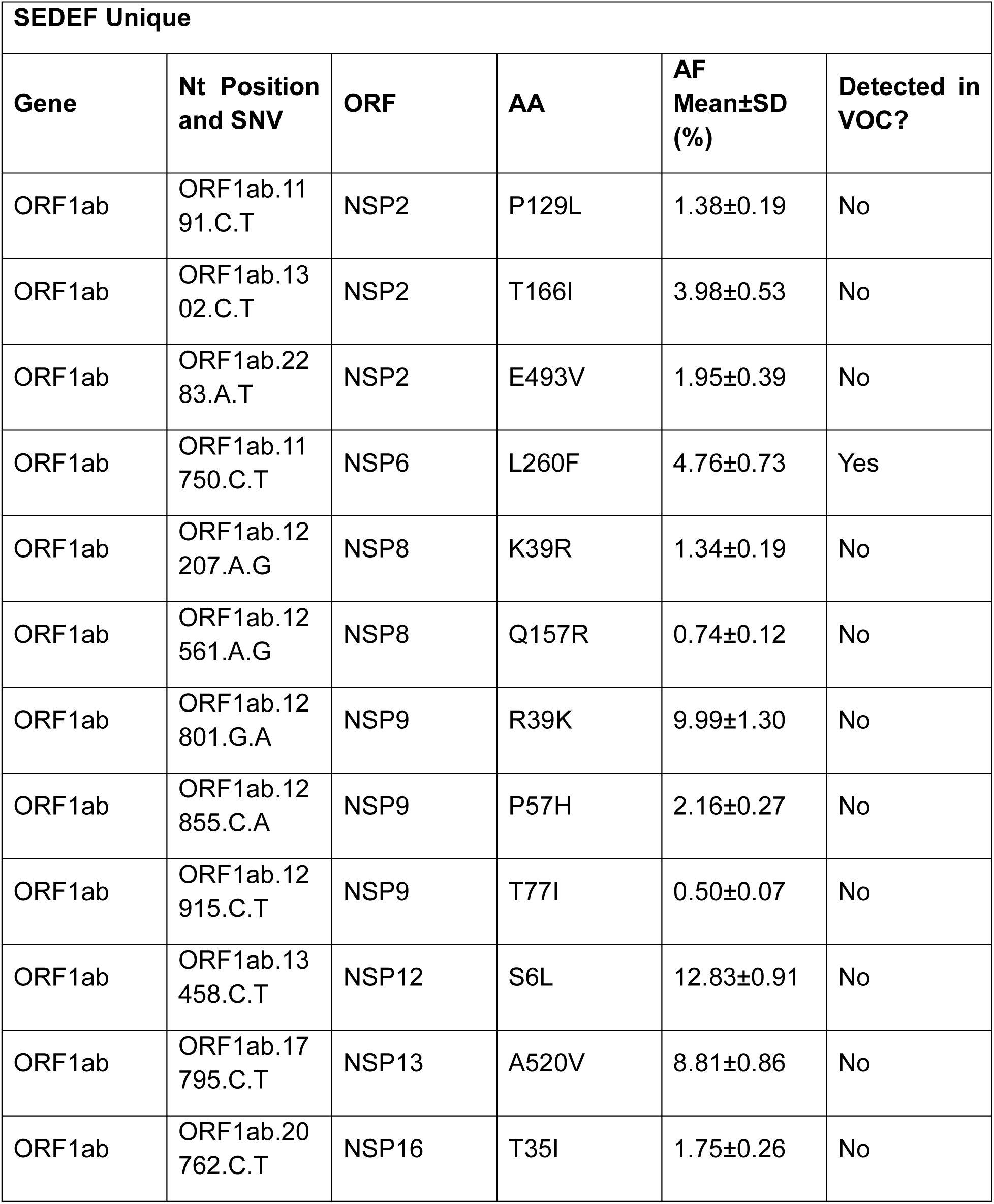
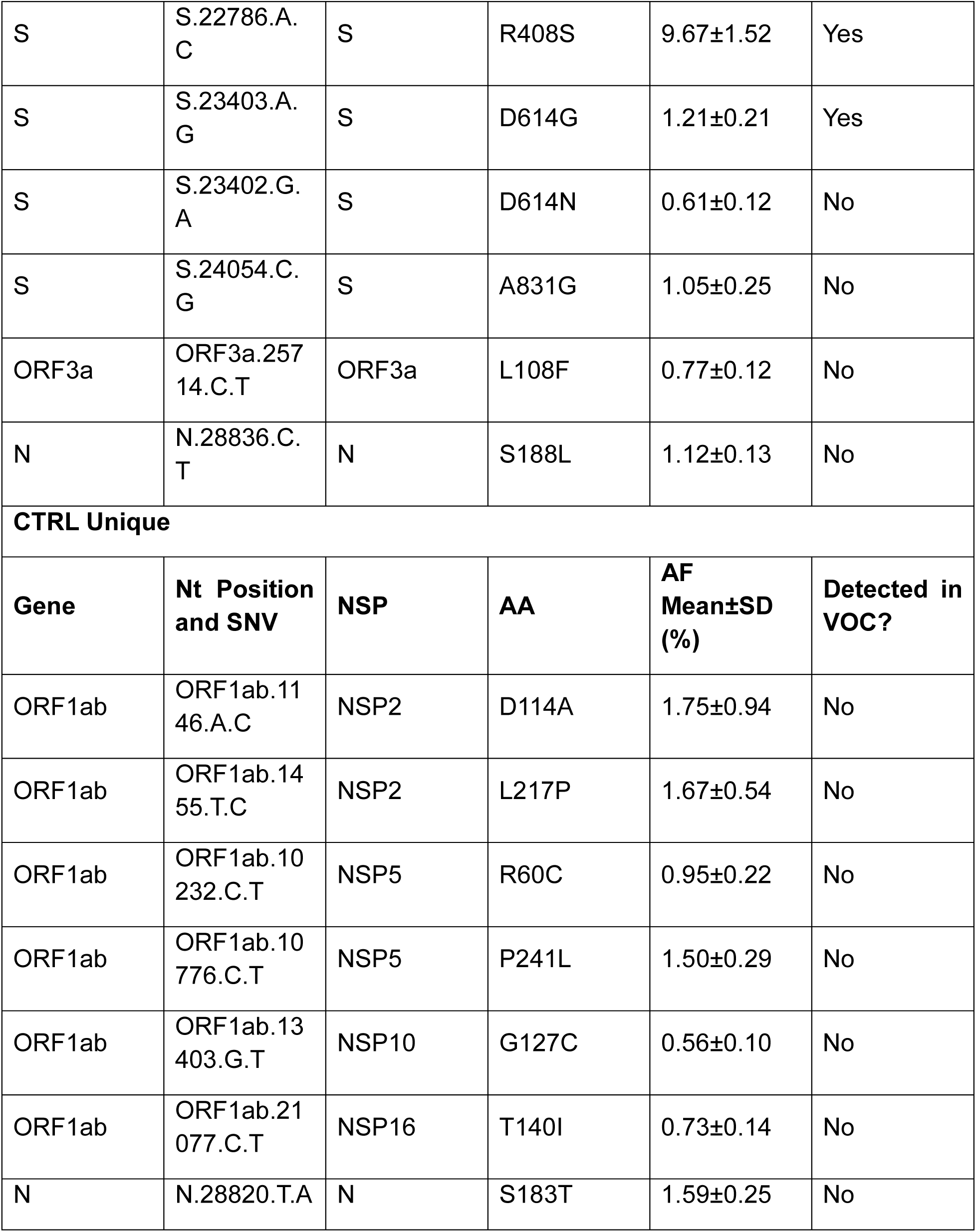
Single nucleotide variations unique to SEDEF or CTRL diet passage. Amino acid position refers to position within the specific ORF. ORF, open reading frame; nt, nucleotide; SNV, single nucleotide variant; AA, amino acid; AF, allele frequency; SD, standard deviation; VOC, variant of concern.

| **SEDEF Unique** |  |  |  |  |  |
| --- | --- | --- | --- | --- | --- |
| **Gene** | **Nt Position and SNV** | **ORF** | **AA** | **AF Mean±SD (%)** | **Detected in VOC?** |
| ORF1ab | ORF1ab.11<br>91.C.T | NSP2 | P129L | 1.38±0.19 | No |
| ORF1ab | ORF1ab.13<br>02.C.T | NSP2 | T166I | 3.98±0.53 | No |
| ORF1ab | ORF1ab.22<br>83.A.T | NSP2 | E493V | 1.95±0.39 | No |
| ORF1ab | ORF1ab.11<br>750.C.T | NSP6 | L260F | 4.76±0.73 | Yes |
| ORF1ab | ORF1ab.12<br>207.A.G | NSP8 | K39R | 1.34±0.19 | No |
| ORF1ab | ORF1ab.12<br>561.A.G | NSP8 | Q157R | 0.74±0.12 | No |
| ORF1ab | ORF1ab.12<br>801.G.A | NSP9 | R39K | 9.99±1.30 | No |
| ORF1ab | ORF1ab.12<br>855.C.A | NSP9 | P57H | 2.16±0.27 | No |
| ORF1ab | ORF1ab.12<br>915.C.T | NSP9 | T77I | 0.50±0.07 | No |
| ORF1ab | ORF1ab.13<br>458.C.T | NSP12 | S6L | 12.83±0.91 | No |
| ORF1ab | ORF1ab.17<br>795.C.T | NSP13 | A520V | 8.81±0.86 | No |
| ORF1ab | ORF1ab.20<br>762.C.T | NSP16 | T35I | 1.75±0.26 | No |

| S | S.22786.A.<br>C | S | R408S | 9.67±1.52 | Yes |
| --- | --- | --- | --- | --- | --- |
| S | S.23403.A.<br>G | S | D614G | 1.21±0.21 | Yes |
| S | S.23402.G.<br>A | S | D614N | 0.61±0.12 | No |
| S | S.24054.C.<br>G | S | A831G | 1.05±0.25 | No |
| ORF3a | ORF3a.257<br>14.C.T | ORF3a | L108F | 0.77±0.12 | No |
| N | N.28836.C.<br>T | N | S188L | 1.12±0.13 | No |
| **CTRL Unique** |  |  |  |  |  |
| **Gene** | **Nt Position**<br>**and SNV** | **NSP** | **AA** | **AF**<br>**Mean±SD**<br>**(%)** | **Detected in**<br>**VOC?** |
| ORF1ab | ORF1ab.11<br>46.A.C | NSP2 | D114A | 1.75±0.94 | No |
| ORF1ab | ORF1ab.14<br>55.T.C | NSP2 | L217P | 1.67±0.54 | No |
| ORF1ab | ORF1ab.10<br>232.C.T | NSP5 | R60C | 0.95±0.22 | No |
| ORF1ab | ORF1ab.10<br>776.C.T | NSP5 | P241L | 1.50±0.29 | No |
| ORF1ab | ORF1ab.13<br>403.G.T | NSP10 | G127C | 0.56±0.10 | No |
| ORF1ab | ORF1ab.21<br>077.C.T | NSP16 | T140I | 0.73±0.14 | No |
| N | N.28820.T.A | N | S183T | 1.59±0.25 | No |

**Figure 1:**
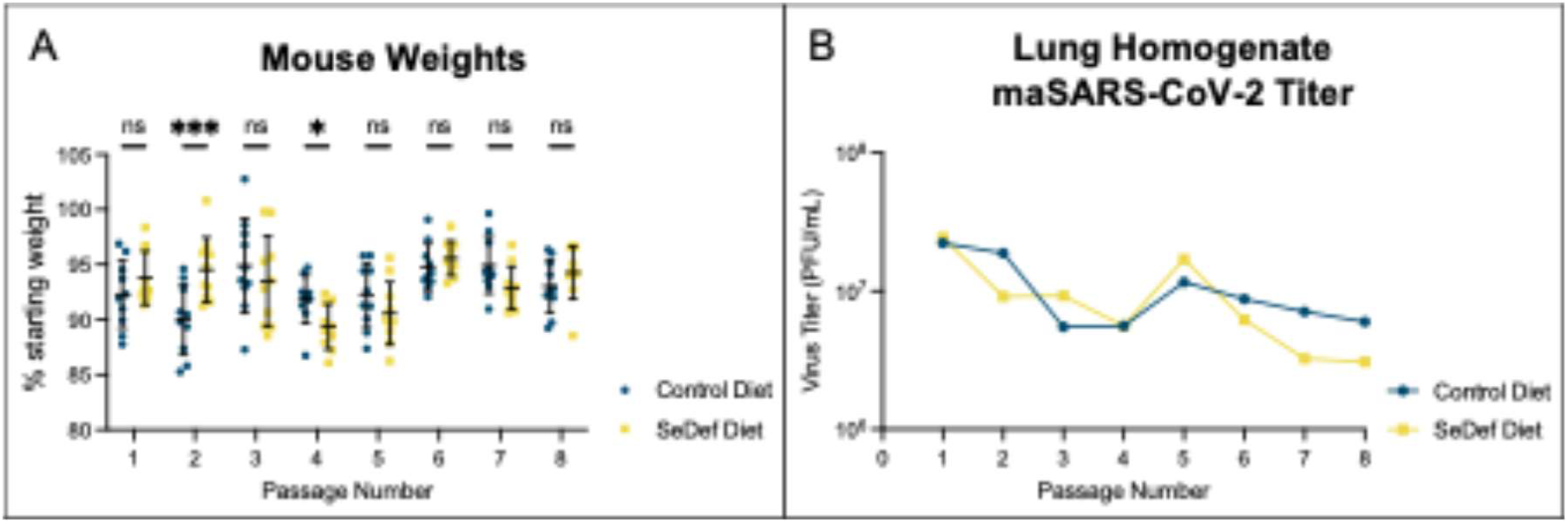
Mouse weight change and lung viral titer during SARS-CoV-2 diet passaging in BALB/C mice. A) Changes in mouse weight two days post viral infection. Data analyzed by twoway ANOVA with Sidak’s multiple comparisons test, comparing control diet to SEDEF diet for each passage. B) Average SARS-CoV-2 viral titer in pooled lung homogenates harvested 2 dpi. ns not significant, *p<0.05, ***p<0.001.

### Deep-sequencing of Passaged SARS-CoV-2 for Variant Analysis

Lung homogenates from the eighth passage cohort of mice were used for Illumina deep sequencing to compare mutational profiles between passage in CTRL and SEDEF backgrounds and to identify emerging, sub-consensus variants. We sequenced individual mouse samples to assess if variation caused by SEDEF passage was unique within an individual mouse or an inter-host effect. Sequencing data was analyzed using the mappgene pipeline, a viral variant calling pipeline for high performance computing platforms^14^. Non-metric multidimensional scaling (NMDS) ordination of single nucleotide variant abundance revealed two visually distinct clusters of sequencing samples (Figure 2A). We observed that samples were clustered based on diet, indicating significant differences in mutant abundance between the CTRL-and SEDEF-passaged virus populations. Analysis for genomic heterogeneity revealed significantly increased positional diversity across nucleotide positions in our SEDEF-passaged population compared to the CTRL-passaged population (Figure 2B).

**Figure 2:**
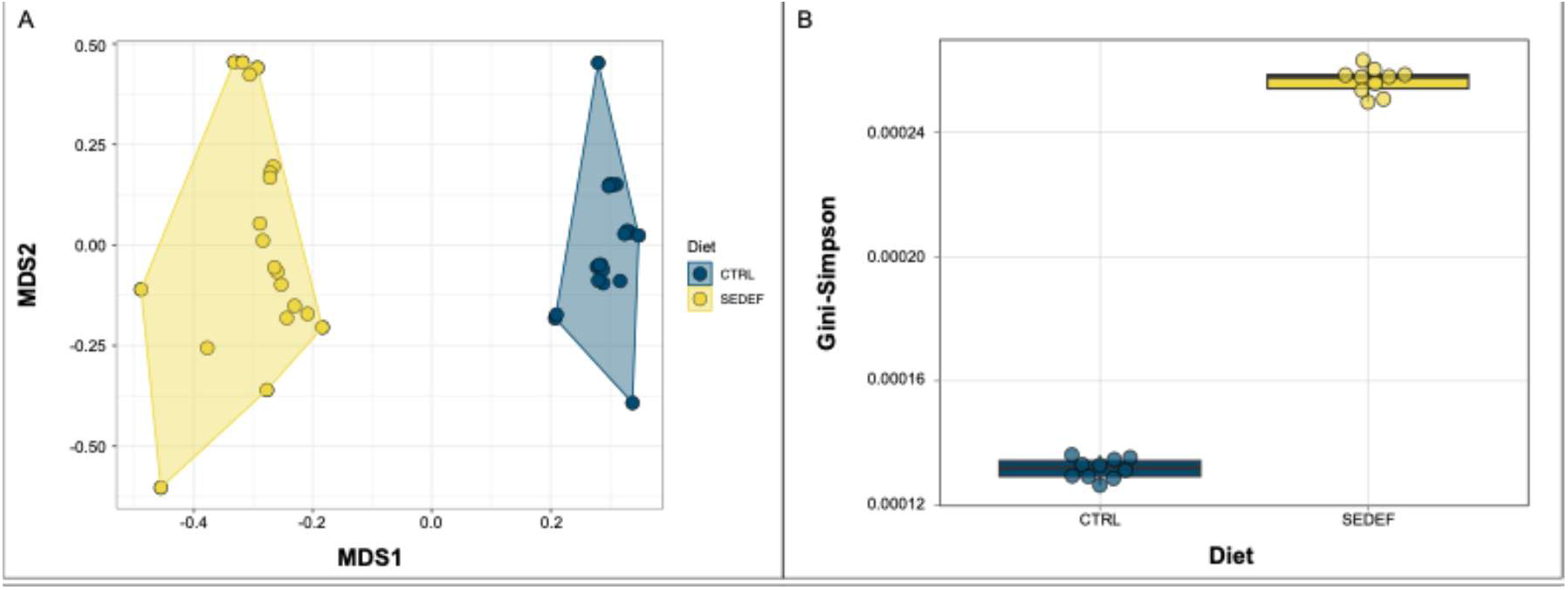
Bioinformatic analysis of mappgene pipeline data. A) NMDS ordination of single nucleotide variant abundance in CTRL and SEDEF virus populations. B) Genomic heterogeneity quantified as average positional diversity per sample using the Gini-Simpson diversity index.

Utilizing our mutant analysis parameters, we found patterns of diet-unique and diet-dominant mutations. We identified 21 diet-unique mutations, 18 of which were unique to SEDEF passage, with the remaining 3 unique to CTRL passage (Table 1). Within each diet-unique group we identified subsets of dominant mutants, defined as mutations observed in both diet groups but only detected above 0.5% allele frequency in one diet. Six SEDEF-unique mutants were identified as SEDEF dominant, while 4 CTRL-unique mutants were identified as CTRL dominant (Supplementary Table 1). To determine if any of our categorized mutants existed in nature as dominant, consensus variants, we searched our down selected mutants through outbreak.info ^15^. Most of our categorized mutants were detected at <0.5% cumulative presence in sequenced global SARS-CoV-2 genomes. None of the mutants categorized as “CTRL Unique” rose above 0.5% cumulative presence in global SARS-CoV-2 sequences or 3% peak frequency (Supplementary Table 2).

Importantly, only viral mutations detected in P8-SEDEF had mutations associated with altered viral fitness and virulence during the COVID19 pandemic. Four of these mutations, (Spike R408S, Spike D614G, NSP2 P129L, NSP6 L260F) are associated with increased disease severity, increased viral transmission, or breakthrough viral infection, and multiple are known mutations in SARS-CoV-2 variant of concern^10,16–24^ (Table 1). Spike R408S, found to interfere with antibody binding, emerged in Omicron clade BA.2 and has been maintained in subsequent Omicron clades^10,16^. R408S was detected at 10% allele frequency in P8-SEDEF samples. Spike D614G arose in early 2020, pre-dating the Alpha variant of concern, and occurred in over 74% of all published SARS-CoV-2 sequences by June 2020^20^. It was the first mutation associated with the global spread of SARS-CoV-2 and increased infectivity of the virus^19,24^. D614G was detected in all P8-SEDEF samples at 1.2% allele frequency. NSP2 P129L is associated with Delta variant lineages and has been identified in breakthrough SARS-CoV-2 infections in Europe, India, and New York^17,21,23^. However, structural modeling suggests P129L would be a destabilizing mutation for NSP2^21^. P129L was detected in all P8-SEDEF sampels at 1.38 allele frequency. NSP6 L260F is associated with increased virulence and mortality in mice, and arose independently across Omicron clades BA.5, BQ.1.1, and XBB.1.16^18,22^. This mutant was detected at 4.8% allele frequency across our sequenced P8-SEDEF samples.

### Characterizing SEDEF-passaged maSARS-COV-2 in Adult BALB/c Mice

Once we obtained our diet passaged virus pools we wanted to understand how maSARS-CoV-2 passaged in a Se deficient diet background changes overall viral severity and mortality. 6–8-week-old, normal diet-fed BALB/c mice were intranasally infected with 10^5^ PFU of P8-CTRL or P8-SEDEF virus survival was tracked for 9 days. Weight loss was not significantly different between P8-CTRL and P8-SEDEF groups (Figure 3A). Fewer P8-SEDEF-infected mice survived compared to P8-CTRL-infected mice, but overall survival was not significantly different between the two groups (Figure 3B). In both virus infection groups, mice that survived infection recovered to or near starting weight approximately 7 dpi. We also investigated *in vivo* viral replication fitness of P8-CTRL and P8-SEDEF virus pools. In serum, P8-CTRL was not detectable above our RT-qPCR limit of detection, whereas P8-SEDEF was detected in most samples (Figure 3C). In lung homogenate, there were no significant changes in genome replication between P8-CTRL and P8-SEDEF virus. The averge genome copy numbers were slightly higher for P8-SEDEF compared to P8-CTRL, but this difference was not significant (Figure 3D). Viral titers trended higher for P8-SEDEF, although differences were not statistically significant (Figure 3E).

**Figure 3:**
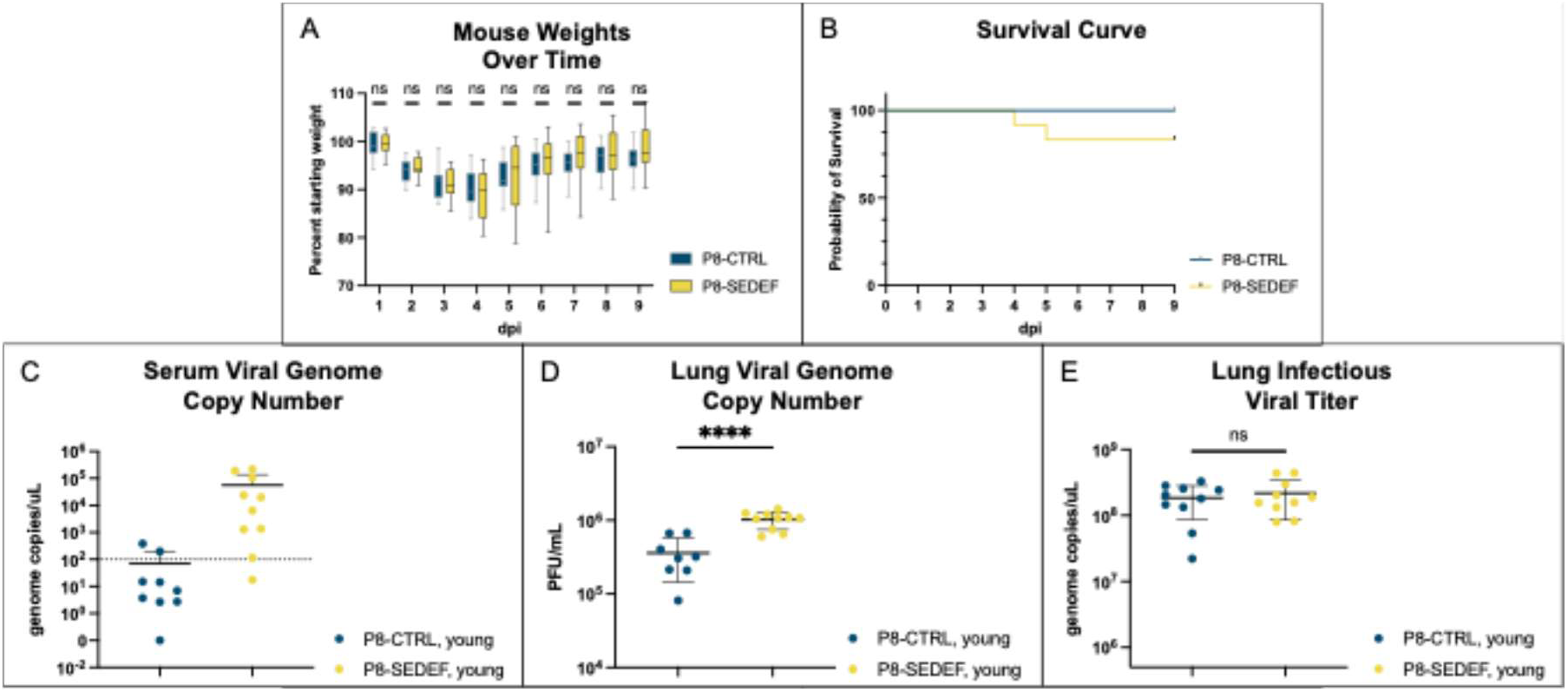
Comparative survival and viral load of adult BALB/C mice following P8-CTRL or P8-SEDEF infection. A) Changes in mouse weight from starting following infection. Analyzed by 2way ANOVA. B) Survival curve following P8-CTRL and P8-SEDEF challenge. C) SARS-CoV-2 genome copy number in lung homogenates 2 dpi measured by RT-qPCR. Dotted line indicates limit of detection D) SARS-CoV-2 genome copy number in serum 2 dpi measured by RT-qPCR. E) SARS-CoV-2 infections viral titer in lung homogenates 2 dpi measure by plaque forming unit assay. ns not significant, **** p<0.0001.

Innate immune responses are a host’s first line of defense against viral infections, including interferon production, cytokine secretion, and downstream immune cell infiltration^10,25^. Cytokine levels were assessed in lung homogenates 2 dpi at peak and near-peak expression following P8-CTRL and P8-SEDEF infection in BALB/c mice^26,27^. For 12 of the 13 cytokines assayed we found P8-SEDEF infection caused significantly increased cytokine expression compared to P8-CTRL infection (Figure 4 A-K, M). The only cytokine with unchanged expression between the two infection groups was CXCL10 (Figure 4L). Both P8-CTRL and P8-SEDEF infection caused increased cytokine expression compared to mock infection, with a mix of significant and non-significant increases.

**Figure 4:**
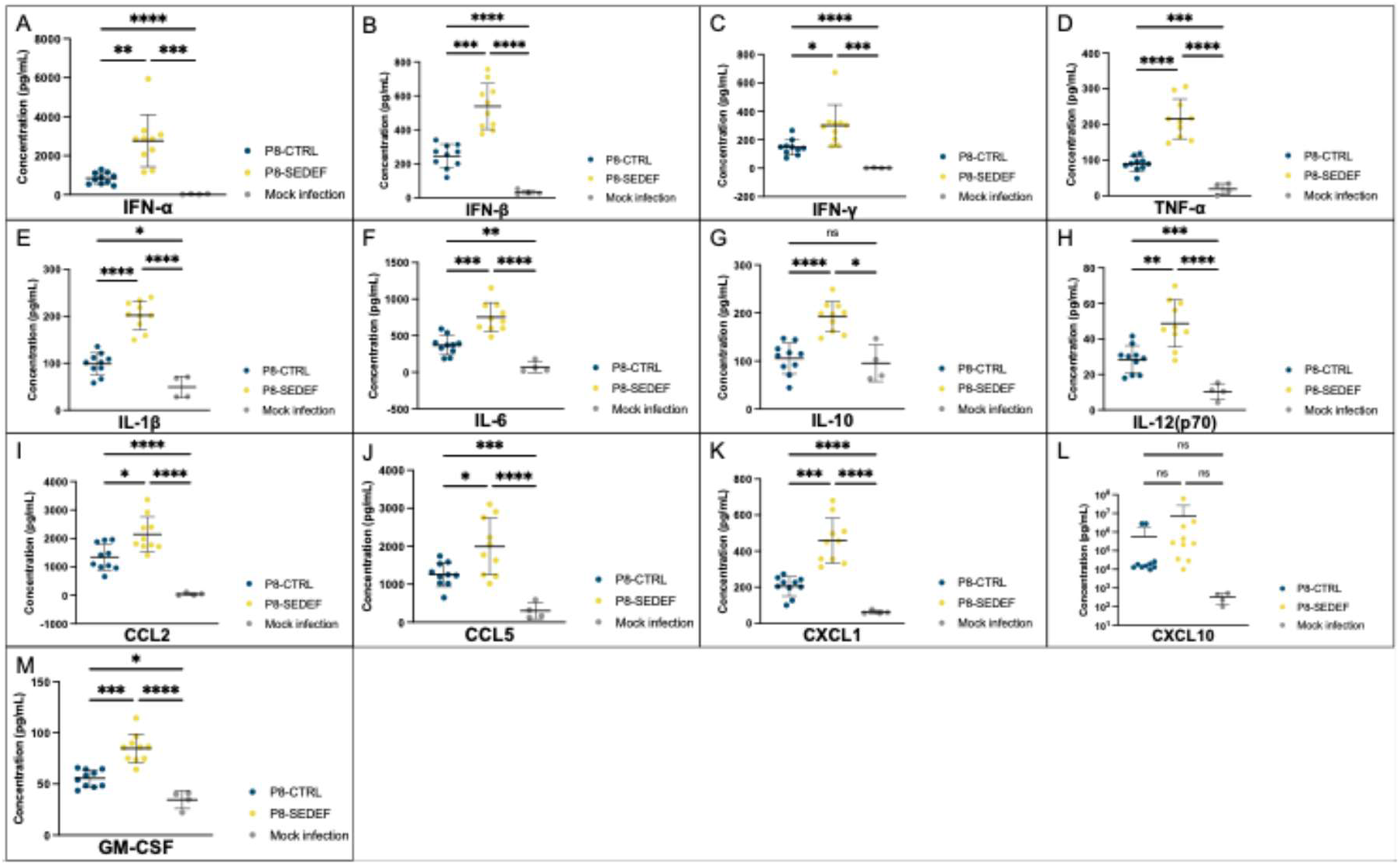
Anti-viral cytokine expression in adult mice lung homogenates following P8-CTRL or P8-SEDEF infection. Cytokine expression measured by LEGENDPlex flow cytometry assay two dpi. A) IFN-⍺, B) IFN-β, C) IFN-Ɣ, D) TNF-⍺, E) IL-1β, F) IL-6, G) IL-10, H) IL-12(p70), I) CCL2, J) CCL5, K) CXCL1, L) CXCL10, and M) GM-CSF. Analyzed by Brown-Forsyth and Welch ANOVA test with Dunnet’s T3 multiple comparisons test. ns, not significant, * p<0.05, ** p<0.01, *** p<0.001, **** p<0.0001.

### Characterizing SEDEF-passaged maSARS-COV-2 in Aged BALB/c Mice

Older adults exhibit dysregulated immune and hemostasis regulation, and these factors contribute to increased susceptibility to infectious diseases like SARS-CoV-2^28,29^. The COVID-19 pandemic revealed a correlation between age and severe disease, with aged populations accounting for the majority of severe COVID-19 cases^29,30^. We hypothesized that aged mice (>10 months old) mice would be more susceptible to changes in disease severity caused by SEDEF passage and could serve as a more sensitive model for observing phenotypic differences between our CTRL-and SEDEF-passaged virus pools. Aged mice were infected with 10^4^ plaque forming units (PFU) of P8-CTRL or P8-SEDEF virus, replicating infectious doses for survival experiments in aged mice conducted in Leist et al., 2020^31^. Weight loss was not significantly different between the two infection groups (Figure 5A). Survival rates were near identical between the two infectioun groups, with over half the mice in both groups succumbing to infection by 4 dpi. The P8-SEDEF group experienced 100% mortality with the P8-CTRL group experience 91% mortality (Figure 5B). Similar survival profiles were also observed following infecting with a log lower dose of virus, with the P8-SEDEF group mice experiencing 100% mortality and the P8-CTRL group experiencing ∼75% mortality (Figure 5C). The mice in this experiment also did not show significant differences in weight loss and recovery (Figure 5D). Viral genome copy numbers in both serum (Figure 5E) and lung homogenates (Figure 5F) did not differ significantly between P8-CTRL and P8-SEDEF infection. In lung homogenates we observed a significant decrease in lung infectious virus in P8-SEDEF infection compared to P8-CTRL infection (Figure 5G).

**Figure 5:**
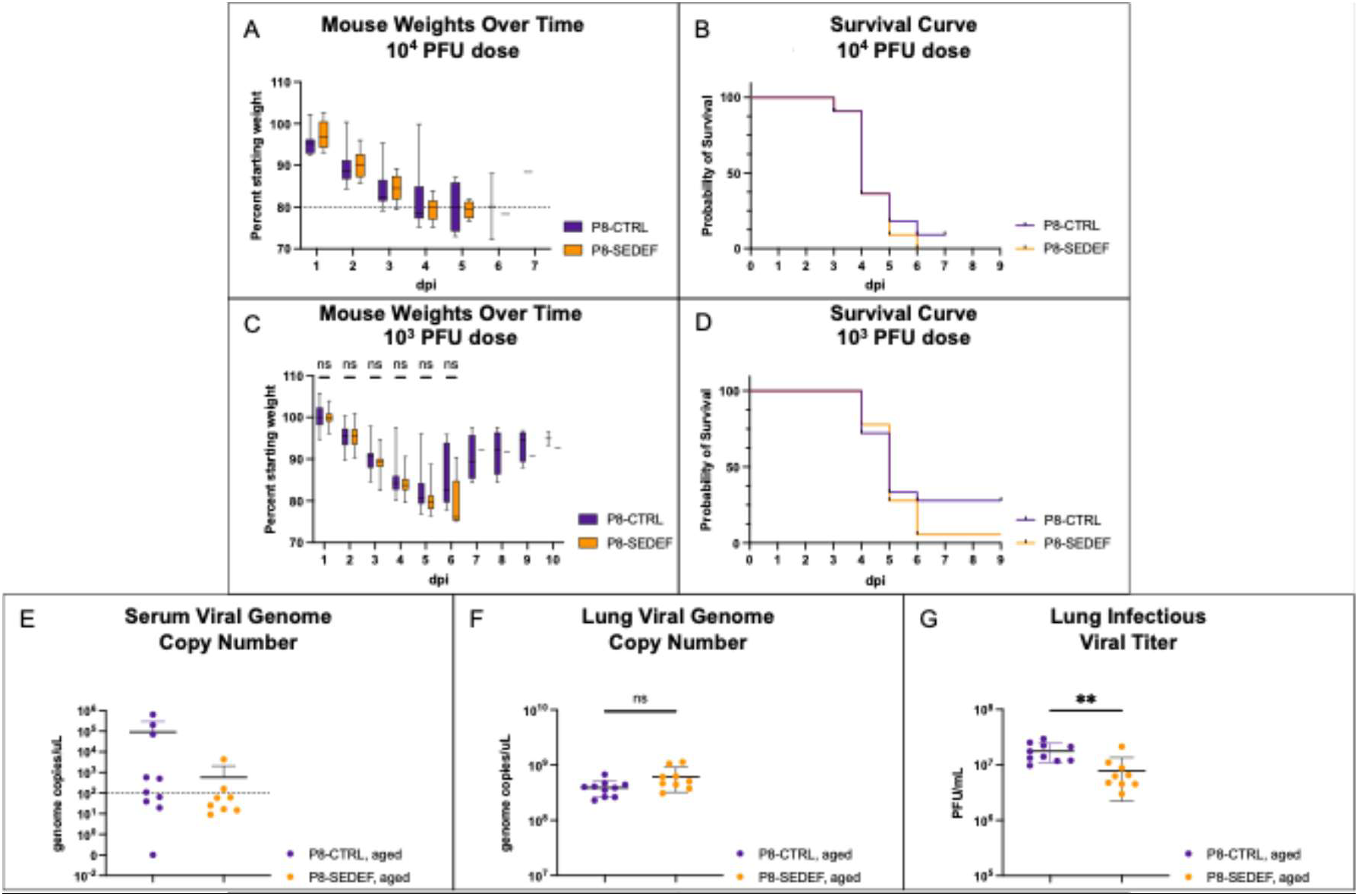
Comparative survival and viral load of aged BALB/C mice following P8-CTRL or P8-SEDEF infection. A) Mouse weights as percentage of starting weight over time following 104 PFU challenge. B) Survival curve following P8-CTRL and P8-SEDEF 104 PFU challenge. C) Mouse weights as percentage of starting weight over time following 103 PFU challenge. D) Survival curve following P8-CTRL and P8-SEDEF 103 PFU challenge. E) SARS-CoV-2 genome copy number in serum 2 dpi measured by RT-qPCR. F) SARS-CoV-2 genome copy number in serum 2 dpi measured by RT-qPCR. G) SARS-CoV-2 infections viral titer in lung homogenates 2 dpi measure by PFU assay. ns not significant; **, p<0.01.

As in our adult mouse cohorts, cytokine levels were assessed in eldery mouse lung homogenates 2 dpi. In general cytokine expression was not significantly different between P8-SEDEF and P8-CTRL infection (Figure 6). IL-1β was the only cytokine with significant expression changes between the two infection groups, showing significantly decreased expression during P8-SEDEF infection compared to P8-CTRL (Figure 6E). IFN-⍺ (Figure 6A), IFN-β (Figure 6B), IFN-Ɣ (Figure 6C), and CCL2 (Figure 6I) were expressed at moderately decreased levels during P8-SEDEF infection, while TNF-⍺ (Figure 6D) and CXCL1 (Figure 6K) were expressed at moderately increased levels in the same mice, all compared to P8-CTRL infection. Both P8-CTRL and P8-SEDEF infection caused increased cytokine expression compared to mock infection with a mix of significant and non-significant increases (Figure 6 A-M). IL-6, CXCL1, and GM-CSF (Figure 6 F, K, and M) expression did not show any notable expression increase from mock infection.

**Figure 6:**
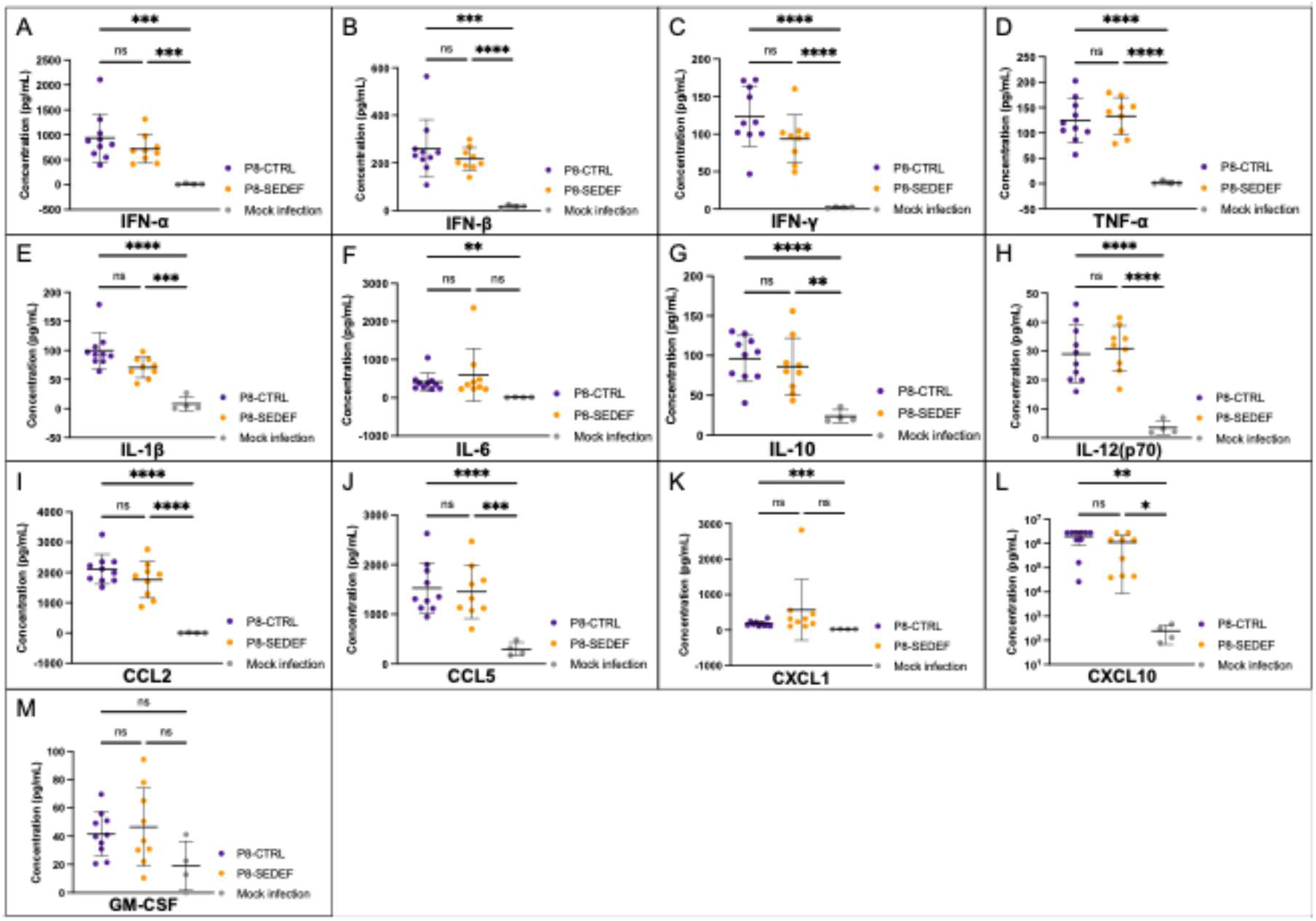
Anti-viral cytokine expression in aged mice lung homogenates following P8-CTRL or P8-SEDEF infection. Cytokine expression measured by LEGENDPlex flow cytometry assay two dpi. A) IFN-⍺, B) IFN-β, C) IFN-Ɣ, D) TNF-⍺, E) IL-1β, F) IL-6, G) IL-10, H) IL-12(p70), I) CCL2, J) CCL5, K) CXCL1, L) CXCL10, and M) GM-CSF. Analyzed by Brown-Forsyth and Welch ANOVA test with Dunnet’s T3 multiple comparisons test. ns, not significant, * p<0.05, ** p<0.01, *** p<0.001, **** p<0.0001.

## Discussion

Individual virions in an RNA virus population do not all carry an identical, consensus genome. Instead, they exist as a large, diverse, intra-host population of variant genomes known as a “quasispecies” which contain sub-consensus mutations^32^. If sub-consensus mutations are beneficial for viral fitness and adaptation, they can rise to become dominant consensus variants or stay maintained at sub-consensus frequency in the quasispecies^13^. Since the beginning of the COVID-19 pandemic numerous SARS-CoV-2 variants emerged and replaced dominant circulating viral strains, with mutations arising from likely a wide range of adaptation events and causes^10,33^. We performed deep sequencing on our CTRL and SEDEF passaged maSARS-CoV-2 populations to determine the following: how does SEDEF impact genomic diversity in maSARS-CoV-2, are these genomic variants consensus or sub-consensus, and can SEDEF produce variants that occurred naturally in SARS-CoV-2?

SEDEF diet passage of ma-SARS-CoV-2 genereated increased single nucleotide variant abundance and genomic heterogeneity in the viral population as compared to CTRL diet passage. We observed an increased number of unique single nucleotide variants in our SEDEF passaged virus compared to our CTRL passaged virus. Mutations identified as unique to SEDEF passage were present as subconsensus rare variants, only detected using ultra-deep sequencing. Our findings are similar to those observed in studies with CVB3. In these studies, researchers found that multiple rounds of CVB3 passage through SEDEF mice produced cardio-virulent mutants that had been observed in CVB3 endemic regions of China^4,9^. One notable difference between our work and the work in CVB3 is observing mutants at the subconsensus versus the consensus level. Our study fouces on the impact of SEDEF passage on expanding the mutant spectra at subconsenses levels, whereas the CVB3 study focused on SEDEF-generated mutations present in the viral genome consensus sequence. Still, these works supports a SEDEF mouse model as a viable tool for identifying putative viral variants that impact virulence and pathogenesis. Further passaging of maSARS-CoV-2 through SEDEF mice could reveal the rise of sub-consensus mutations to consensus sequence, and could allow for optimization of the SEDEF mouse model as a pipeline protocol for future uses.

We infected normal diet-fed mice with P8-CTRL and P8-SEDEF virus pools to understand how the increased mutant spectrum of P8-SEDEF impacts *in vivo* viral infection. In adult and aged mice, P8-SEDEF infection trended to decreased survival over P8-CTRL infection, with weight change between the two viruses not differing in both age groups. However, these data did not support our hypothesis that SEDEF-passage would significantly impact viral disease severity. Both ages groups did not show significant differences in serum genome copy number between the two viruses. Significant differences in viral replication between P8-CTRL and P8-SEDEF in the lungs were observed in both age groups but via different metrics. Adult mice showed differences in P8-CTRL and P8-SEDEF genome copy number, while aged mice showed differences between P8-CTRL and P8-SEDEF infectious titers. Previous studies investigating RNA viruses and Se deficiency have found SEDEF-passaged virus demonstrated significant changes to replication and lethality upon challenge in a normal diet-fed mice^8,9,34–36^, but differences in methods between this study and previous could explain these differences. One study investigating Sendai virus passage in a Se deficient mouse model found that SEDEF-passage virus caused significantly increased weight loss and significantly reduced viral titers in the lungs^34^. Previously mentioned studies with CVB3 found that SEDEF-passaged virus produced significantly higher virus titers than control diet-passaged virus in the hearts of normal diet-fed mice^4,9^. The differences between our data and previous studies’ data could likely be attributed to the different viruses being studied. Another explanation for our observations could be the nature of our mutant P8-SEDEF virus population. As discussed, the increased genomic heterogeneity we observed in our P8-SEDEF population was far below consensus level. We hypothesize that variants present in P8-SEDEF that significantly impact viral lethality and replication do not exist at a high enough allele frequency to have significant impacts on these aspects of SARS-CoV-2 infection.

A major hallmark of severe COVID-19 disease is the development of an exaggerated acute secretion of pro-inflammatory cytokines and systemic hyperinflation called a cytokine storm^25^. SARS-CoV-2 variants of concern, including Alpha and Omicron variants, have evolved to further aggravate the cytokine storm during COVID-19^10^. We measured anti-viral cytokine production following infection with our P8-CTRL and P8-SEDEF virus pools to determine how SEDEF-passaged virus impacts host innate immune responses. In a panel of 13 antiviral cytokines, we observed variable changes in cytokine expression between age groups. In adult mice, P8-SEDEF infection caused significantly increased cytokine expression for 12 of the measured cytokines compared to P8-CTRL infection. CXCL10 was the single cytokine with significantly unchanged expression between P8-CTRL and P8-SEDEF infection in the adult mice (Figure 4). COVID-19 severity increases with age, with aged adults account for the largest percentage of severe COVID-19 cases^29,37^. Increased basal IL-6 and TNF-⍺ expression in aged adults is associated with increased infection susceptibility and is thought to contribute to hyperimmune activation seen in older adults with COVID-19^28,37^. In aged mice we curiously only observed significant changes in IL-1β expression between P8-SEDEF and P8-CTRL infection groups and did not observe significant changes in the remaining assayed cytokines (Figure 6). Our data show that adult mice are more susceptible to immune changes caused by increased viral diversity than aged mice. It is possible that virus-induced cytokine expression is not altered at early time points during infection due to increased basal inflammatory cytokine expression associated with age. Future studies may assay cytokine expression at later time points in aged mice to determine if increased cytokine expression, like that seen in adult mice, would occure in aged mice cohorts.

Undernutrition, particularly Se deficiency, has been recognized as a critical factor that compromises host immune responses and increases infection susceptibility^38,39^. A growing body of ecological and epidemiological evidence indicates that Se deficiency increases the severity and mortality of various viruses including hantavirus, HIV, influenza, and SARS-CoV-2^4,9,38,40–44^. Despite these important associations, there is lacking data on how Se deficiency drives emergence of more virulent viral genotypes and phenotypes. This study supports our hypothesis that passage through a SEDEF population increases viral mutant diversity, and can contribute to variant emergence.The consistent production of mutants that arose naturally since human SARS-CoV-2 emergence demonstrates the utility of a SEDEF animal model as a method to forecast putative emerging viral variants in global regions of Se-deficiency. The detection of these mutations by deep sequencing and the mappgene pipeline highlights the utility of theses tools for detecting emergent viral variants in host populations. Our studies sequenced maSARS-CoV-2 at a single time point following multiple passages through SEDEF mice, and sequencing of maSARS-CoV-2 after each mouse passage could reveal more information about variant emergence and dynamics in the viral population. While downstream investigation revealed limited alterations to survival or viral replication, SEDEF passage did cause significant alterations in host immune response to infection. Together, our data demonstrate host Se-deficiency as a driver of viral populaiton diversification, and that RNA virus passage through *in vivo* Se-deficiency models could be a critical tool for identifying putative variants in pandemic-potential viruses in Se-deficient regions.

## Materials and Methods

### Cells and Viruses

VeroE6 and VeroE6-TMPRSS2 cells were maintained in 1X DMEM medium with 10% FBS, 1% HEPES, 1% sodium pyruvate, 1% non-essential amino acids, and 1% penicillin-streptomycin. Cells were cultured in a 37°C incubator with 5% CO_2_. Mouse-adapted MA 10 Variant infectious clone (ic2019-nCov MA10) was obtained from BEI Resources (NR-55329) and referred to in this text as “maSARS-CoV-2”. For stock virus generation, 10^6^ VeroE6-TMPRSS2 cells were infected with 20 µL of BEI stock virus and supernatant was harvested 3 days post infection (dpi). 10 µL of VeroE6-TMPRSS2-passaged virus was used to inoculate 10^6^ Vero-TMPRSS cells and supernatant harvested 3 dpi. The titer of the Vero-TMPRSS passaged virus was quantified by plaque forming unit (PFU) assay, described below.

### Plaque Forming Unit Assay

MaSARS-CoV-2 samples were diluted ten-fold in serum-free culture media (1X DMEM, 1% each HEPES, non-essential amino acids, sodium pyruvate, and penicillin-streptomycin). In 6-or 12-well plates, 200 µL of virus dilutions were used to inoculate confluent VeroE6-TMPRSS2 cells. Cells were inoculated for 30 minutes at 37°C, with plate rocking every 5 minutes. Inoculum was overlayed with a 1:1 ratio of 2.4% microcrystalline methylcellulose in water and plaque assay overlay medium (2X DMEM, 8% FBS, and 2% each HEPES, non-essential amino acids, sodium pyruvate, and penicillin-streptomycin). 2 mL of overlay was used for 6-well plates, and 1 mL overlay was used for 12-well plates. 72 hours post infection, overlay was removed, and wells were washed with 1X PBS. Wells were stained with crystal violet in 70% methanol for 15 minutes.

### Biosafety Methods

All infectious maSARS-CoV-2 work was reviewed and approved by the LLNL Institutional Biosafety Committee and was conducted in LLNL’s BSL-3 laboratories following established BSL-3 standard operating procedures. The LLNL Institutional Animal Care and Use Committee (IACUC) reviewed and approved all animal experiments, which were performed in an AAALAC International-accredited facility. maSARS-CoV-2 viral RNA was extracted from lung homogenate and serum using the QIAgen Viral RNA extraction kit in accordance with in-house validated inactivation procedures verified by viability testing prior to removal from the BSL-3 Facility for downstream analysis in BSL-2 laboratories.

### Animal Ethics Statement

These studies were carried out in strict accordance with the recommendations in the Guide for the Care and Use of Laboratory Animals and the National Institute of Health. All efforts were made to minimize suffering of animals. All animals were housed in ABSL-3 conditions in an AAALAC International-accredited facility, and the protocol was approved by the LLNL Institutional Animal Care and Use Committee (IACUC; Protocol 315, approved October 17, 2022), which includes ethics in evaluation of protocols.

### Mouse maSARS-CoV-2 Passaging and Survival Studies

BALB/c mice were fed control or selenium-deficient chow for 4 weeks prior to infection (Table 2). For the first passage, ten mice in both diet groups were intranasally infected with 10^5^ PFUs of parental maSARS-CoV-2. Two days post-infection (dpi), mice were euthanized and total lungs were harvested from each mouse. Lungs were homogenized in 1 mL PBS in 2 mL bead ruptor pre-filled bead tubes (Omni International) with disruption at setting 5 for 20 seconds using a Bead Ruptor 4 (revvity). Disruption was repeated until no solid lung tissue was observed. Lung homogenates were centrifuged at 10,000 rcf for 5 minutes. Clarified lung homogenate supernatant was transferred to a 15 mL conical tube, and individual samples were pooled within each diet group. An aliquot of pooled homogenate for each diet group was used to determine the virus titer by PFU assay.

**Table 2:** Composition of experimental diets. Diets were commercially produced by Teklad Custom Diets, Madison, WI.

| TD.96363 - Control Diet |  |
| --- | --- |
| Formula | g/kg |
| Torula Yeast | 300.0 |
| DL-Methionine | 3.0 |
| Sucrose | 591.0 |
| Corn Oil | 50.0 |
| Mineral Mix, AIN-76 (170925) | 35.0 |
| Calcium Carbonate | 11.0 |
| Vitamin Mix, Teklad (40060) | 10.0 |
| TD.92163 - Selenium Deficient Diet |  |
| Formula | g/kg |
| Torula Yeast | 300.0 |
| DL-Methionine | 3.0 |
| Sucrose | 591.0 |
| Corn Oil | 50.0 |
| Mineral Mix, Selenium Deficient (80313) | 35.0 |
| Calcium Carbonate | 11.0 |
| Vitamin Mix, Teklad (40060) | 10.0 |

For subsequent mouse passages, pooled lung homogenate was used to infect a new cohort of on-diet BALB/c mice. Control diet-passaged homogenate was used to infect control diet-fed mice, and selenium deficient-passaged homogenate was used to infected selenium deficient diet-fed mice. The same lung isolation, homogenization, pooling, and tittering method was used for each mouse passage. This process of control-passage and selenium deficient-passage was repeated eight times. Each passage cohort has five males and five females for each diet group. Lung homogenate used for downstream experimental analysis will be referred to by passage number 8 (P8), and from control diet passaging (CTRL) or selenium deficient diet passaging (SEDEF) in this text.

For the remaining mouse studies, mice were anaesthetized using 4-5% isoflurane in 100% oxygen. Mice were inoculated intranasally with 10^5^ PFU (adult) or 10^4^ or 10^3^ PFU (aged) of P8-CTRL or P8-SEDEF pooled lung homogenate. Mock infected mice were inoculated with an equal volume of 1X PBS. Mice were monitored daily for weight change and clinical disease presentation. In our survival experiments, mice were euthanized when dropping below 80% of initial starting weight. For non-survival experiments, mice were euthanized at 2 dpi.

### Mouse Sample Collection

Blood was harvested using cardiac stick and transferred to a 1.5 mL microcentrifuge tube. All blood samples were incubated at room temperature for 30 minutes for coagulation then centrifuged at 2000 rcf for 10 minutes at 4°C to separate blood clot from serum. Serum was transferred by pipetting to a screw-cap freezer vial and stored at -80°C until later analysis. Both lung lobes were harvested from each mouse and cut into smaller pieces with a razor blade. Lung pieces were homogenized as described above. Clarified homogenate was aliquoted into screw-cap freezer vials and stored at -80°C until later analysis.

### ARTIC RT-PCR of maSARS-CoV-2 Mouse Samples

Viral genomic RNA was isolated from unpooled P8 lung homogenate samples using the QIAmp Viral RNA Mini Kit (Qiagen), following the manufacturer’s protocol. Viral RNA was reverse transcribed and PCR (RT-PCR) amplified using the NEBNext® ARTIC SARS-CoV-2 RT-PCR Module kit (New England Biolabs). 2.5×10^7^ genome copies were used as template for reverse transcription, and all samples were amplified using Primer Mix 1 and Primer Mix 2 in separate reactions. All viral RNA samples were RT-PCR amplified in duplicate for both primer mixes. Amplicon size was validated by running amplified samples on a E-Gel 2% agarose gel with SYBR™ Safe (Invitrogen) using a E-Gel Power Snap Electrophoresis system (Invitrogen).

Library preparation was done according to Avila-Herrera et al., 2024 with the following modifications^45^. A total of 40 µL of pooled Primer Pool 1 and Primer Pool 2 ARTIC amplicons were purified by adding 0.8X Ampure XP Magnetic Beads (Beckman Coulter) to each pooled amplicon sample. PCR products were purified following manufacturer’s protocol and resuspended in 22 µL of water. Purified PCR products were quantified using a Qubit fluorimeter (ThermoFisher). 100 ng of each purified PCR sample was used as input for the Illumina DNA Prep Library Kit (Illumina), and library prep was done using the small amplicon manufacturer’s protocol. Completed libraries were quantified and verified on a TapeStation 4200 instrument with appropriate reagents (Agilent Technologies). Libraries were diluted to 4 nM, pooled, and further diluted to a loading concentration of 8 pM. Libraries were sequenced using an Illumina NexSeq 2000 platform using a P1 300 cycle (2 x 150 bp) kit with onboard denaturation and dilution. Raw sequencing data has been deposited in the NCBI Sequence Read Archive under BioProject PRJNA1374194.

### Bioinformatics Analysis

Variants were called using mappgene (https://github.com/LLNL/mappgene) a viral variant calling pipeline for high performance computing environments^14^. Mappgene employs BWA-MEM for read alignment, Ivar 1.4.3 for read trimming, filtering, and variant calling, and LoFreq v2.1.5 for variant calling^46,47^.

### Nucleotide Diversity and Genomic Heterogeneity Analysis

Positional nucleotide diversity was quantified as the Gini-Simpson diversity index of the distribution of nucleotides at each variant locus for each sample. The Gini-Simpson index represents the probability that two randomly chosen bases are different. The Gini-Simpson diversity index is known by many names and is related to Nei and Li’s nucleotide diversity (π), Renyi entropy of order 2, Hill’s effective number of species (N2), and the Simpson index by simple transformations:

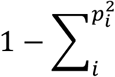

where *p_i_* is the proportion of nucleotide base *i* ^48–50^. Genomic heterogeneity was calculated as the average positional diversity for each sample.

### Mutant Analysis

Mutants were chosen for analysis if they were detected in a least 9 out of 10 mice per sex for each diet group at an allele frequency >0.5% as detected by Lofreq. Mutants were categorized as “diet unique” if they were detected only in a single diet group. Mutants were categorized as “diet dominant” if they were detected in both diet groups but only detected at an allele frequency >0.5% in one. All categorized mutants were searched on outbreak.info (https://outbreak.info/) for detection and prevalence in globally sequenced SARS-CoV-2 sequences^15,51^. Outbreak.info was access on 11 October 2025. On accession date, all data were as of March 5, 2025, 10:41 AM. Data taken for “Worldwide” location search setting^15^.

### Quantification of maSARS-CoV-2 Genomic RNA by RT-qPCR

Viral genomic RNA was isolated from viral stock, mouse lung homogenate, and mouse serum samples using the QIAmp Viral RNA Mini Kit (Qiagen), following the manufacturer’s protocol. Viral genomic RNA was quantified using the PrimeTime One-Step RT-qPCR Master Mix (IDT) using 2 µL of RNA as template. IDT primers and probes RdRP_SARSr_F2 Forward Primer (10006860), RdRP_SARSr_R1 Reverse Primer (10006881), and RdRP_ SARSr_P1 (FAM) Probe (10006884) were used to amplify maSARS-CoV-2 genome, with the IDT 2019-nCoV_RdRP (ORF1ab) Positive Control (10006897) used as a standard curve to quantify genomic copy numbers. RNA samples were run in duplicate.

### Cytokine Analysis

Lung homogenates were harvested from mice 2 dpi and analyzed using the LEGENDPlex™ Mouse Anti-Virus Response Panel (13-plex, V-bottom plate, Biolegend) according to manufacturer’s protocol. Samples were assessed using an Accuri™ B6 flow cytometer (Becton Dickinson). Data were analyzed using the online Data Analysis Software Suite for LEGENDPlex™ (BioLegend, https://www.biolegend.com/en-ie/immunoassays/legendplex/support/software), and statistical analysis was performed using GraphPad Prism.

### Use of Artificial Intelligence (AI)-Assistend Tools

The authors used LivChat, a LLNL generative AI tool, to assist with drafting this manuscript’s abstract and identification of references relevant to the cytokines analyzed in this work. No AI tool was used to generate, analyze, or interpret experimental data. All AI-assisted content was reviewed, edited, and validated by the authors. The authors assume full responsibility for the content and compliance of the submitted work.

## Acknowledgements

This work was performed under the auspices of the U.S. Department of Energy by Lawrence Livermore National Laboratory under Contract DE-AC52-07NA27344 and was supported by the Lawrence Livermore National Laboratory-Laboratory Directed Research and Development Program under Project No. 49843/23-ERD-012. The following reagents were obtained through BEI Resources: NR-55329 Mouse-adapted MA 10 Variant (in isolate USA-WA1/2020 backbone), Infectious Clone (ic2019-nCoV MA10).

## Data Availability

Raw sequencing data has been deposited in the NCBI Sequence Read Archive under BioProject accession number PRJNA1374194.

---

**Supplementary Table 1:** Single nucleotide variations that were observed in both diet groups but more dominantly in one. Variants that were observed above 0.5% allele frequency in one diet, and below 0.5% allele frequency in the other diet. ORF, open reading frame; nt, nucleotide; SNV, single nucleotide variant; AA, amino acid; AF, allele frequency; SD, standard deviation.

| **SEDEF Dominant** |  |  |  |  |  |  |
| --- | --- | --- | --- | --- | --- | --- |
| **Gene** | **Nt Position and SNV** | **ORF** | **AA** | **Diet** | **Mouse Count** | **AF Mean±SD (%)** |
| ORF1ab | ORF1ab.1191.C.T | NSP2 | P129L | CTRL | 19 | 0.32±0.09 |
|  |  |  |  | SEDEF | 20 | 1.38±0.19 |
| ORF1ab | ORF1ab.11750.C.T | NSP6 | L260F | CTRL | 16 | 0.25±0.07 |
|  |  |  |  | SEDEF | 20 | 4.76±0.73 |
| ORF1ab | ORF1ab.12801.G.A | NSP9 | R39K | CTRL | 3 | 0.24±0.04 |
|  |  |  |  | SEDEF | 20 | 9.99±1.30 |
| S | S.22786.A.C | S | R408S | CTRL | 15 | 0.33±0.04 |
|  |  |  |  | SEDEF | 20 | 9.67±1.52 |
| S | S.23402.G.A | S | D614N | CTRL | 20 | 0.35±0.11 |
|  |  |  |  | SEDEF | 20 | 0.61±0.12 |
| ORF3a | ORF3a.25714.C.T | ORF3a | L108F | CTRL | 1 | 0.22±N/A |
|  |  |  |  | SEDEF | 20 | 0.77±0.12 |
| **CTRL Dominant** |  |  |  |  |  |  |
| **Gene** | **Nt Position and SNV** | **ORF** | **AA** | **DIET** | **Mouse Count** | **AF Mean±SD (%)** |
| ORF1ab | ORF1ab.1146.A.C | NSP2 | D114A | CTRL | 20 | 1.75±0.94 |
|  |  |  |  | SEDEF | 20 | 0.47±0.17 |
| ORF1ab | ORF1ab.10232.C.T | NSP5 | R60C | CTRL | 20 | 0.95±0.22 |
|  |  |  |  | SEDEF | 8 | 0.25±0.02 |
| ORF1ab | ORF1ab.10776.C.T | NSP5 | P241L | CTRL | 20 | 1.50±0.29 |
|  |  |  |  | SEDEF | 20 | 0.35±0.08 |
| ORF1ab | ORF1ab.21077.C.T | NSP16 | T140I | CTRL | 19 | 0.73±0.14 |
|  |  |  |  | SEDEF | 1 | 0.31±N/A |

**Supplementary Table 2:** Information for detection and presence in surveillance data sourced from outbreak.info. “Total Detections” represents worldwide number of reported sequences containing the mutation. “Cumulative Prevalence” represents the apparent ratio of the sequences containing a mutant to all sequences collected since the identification of that mutant worldwide. “First Detected” and “Last Detected” dates based on the sample collection dates. “Peak Frequency” represents the average daily prevalence worldwide of the analyzed mutant based on reported sample collection dates. *, mutants found in VOCs. ^†^, mutants categorized as diet dominant.

| **SEDEF** |  |  |  |  |  |
| --- | --- | --- | --- | --- | --- |
| **Mutant** | **Total Detections** | **Cumulative Presence** | **First Detected** | **Last Detected** | **Countries Detected** |
| NSP2.P129L<sup>†</sup> | 134794 | 1% | 11-Mar-20 | 23-Nov-24 | 172 |
| NSP2.T166I | 10830 | <0.5% | 02-Mar-20 | 18-Nov-24 | 107 |
| NSP2.E493V | 12 | <0.5% | 28-Apr-21 | 11-Sep-23 | 4 |
| NSP6.L260F*<br><sup>†</sup> | 662676 | 4% | 08-Feb-20 | 16-Oct-24 | 189 |
| NSP8.K39R | 441 | <0.5% | 18-Mar | 17-Jun-24 | 38 |
| NSP8.Q157R | 1 | <0.5% | 20-Mar-21 | 20-Mar-21 | 1 |
| NSP9.R39K<sup>†</sup> | 51595 | <0.5% | 05-Apr-20 | 14-Oct-24 | 78 |
| NSP9.P57H | 413 | <0.5% | 30-Mar-20 | 05-Sep-24 | 46 |
| NSP9.T77I | 5894 | <0.5% | 05-Mar-20 | 17-Oct-24 | 89 |
| NSP12.S6L | 21506 | <0.5% | 12-Mar-20 | 21-Oct-24 | 128 |
| NSP13.A520V | 5575 | <0.5% | 09-Apr-20 | 14-Oct-24 | 89 |
| NSP16.T35I | 39784 | <0.5% | 17-Feb-20 | 20-Oct-24 | 129 |
| S.R408S*<sup>†</sup> | 5665386 | 37% | 25-Sep-20 | 10-Dec-24 | 191 |
| S.D614G* | 15421541 | 100% | 15-Jan-20 | 10-Dec-24 | 212 |
| S.D614N<sup>†</sup> | 357 | <0.5% | 24-Mar-20 | 15-Jul-21 | 27 |
| S.A831G | 46 | <0.5% | 26-Jun-21 | 24-Jun-24 | 9 |
| ORF3a.L108F | 31497 | <0.5% | 14-Mar-20 | 19-Oct-24 | 132 |
| N.S188L | 1304 | <0.5% | 06-Mar-20 | 02-Mar-24 | 31 |
| **CTRL** |  |  |  |  |  |

| **Mutant** | **Total**<br>**Detections** | **Cumulative**<br>**Presence** | **First**<br>**Detected** | **Last**<br>**Detected** | **Countries**<br>**Detected** |
| --- | --- | --- | --- | --- | --- |
| NSP2.D114A<sup>†</sup> | 219 | <0.5% | 09-Sep-20 | 08-Jul-24 | 28 |
| NSP2.L217P | 1116 | <0.5% | 20-Mar-20 | 30-Aug-24 | 47 |
| NSP5.R60C<sup>†</sup> | 1565 | <0.5% | 22-Jan-20 | 31-Jul-24 | 61 |
| NSP5.P241L<sup>†</sup> | 10757 | <0.5% | 07-Apr-20 | 18-Oct-24 | 110 |
| NSP10.G127<br>C | 493 | <0.5% | 25-Mar-20 | 24-Jul-24 | 38 |
| NSP16.T140I<sup>†</sup> | 76174 | <0.5% | 09-Mar-20 | 16-Oct-24 | 154 |
| N.S183T | 6024 | <0.5% | 13-Apr-20 | 21-Aug-24 | 48 |

